# Regulation of cGAS activity through RNA-mediated phase separation

**DOI:** 10.1101/2020.09.27.316166

**Authors:** Silian Chen, Miao Rong, Yun Lv, Deyu Zhu, Ye Xiang

## Abstract

Cyclic GMP-AMP synthase (cGAS) is a double-stranded DNA (dsDNA) sensor that functions in the innate immune system. Upon binding dsDNA in the cytoplasm, cGAS and dsDNA form phase-separated aggregates in which cGAS catalyzes synthesis of 2’3’-cyclic GMP-AMP that subsequently triggers a STING-dependent, type I IFN response. Here, we showed that cytoplasmic RNAs, especially tRNAs, regulate cGAS activity. We discovered that RNAs did not activate cGAS but rather promoted phase separation *in vitro*. In cells, cGAS colocalized with RNAs and formed phase-separated granules even in the absence of cytoplasmic dsDNA. An Opti-prep gradient analysis of cell lysates showed that the endogenous cGAS was associated with cytoplasmic RNAs in an aggregative form. Further *in vitro* assays showed that RNAs compete for binding of cGAS with dsDNA and inhibit cGAS activity when the dsDNA concentration is high and promote the formation of phase separations and enhance cGAS activity when the dsDNA concentration is low. Thus, cytoplasmic RNAs regulate cGAS activity by interfering with formation of cGAS-containing aggregates.

## Introduction

Recognition of pathogen-derived nucleic acids by protein sensors allows the innate immune system to sense infection and initiate host defense mechanisms (*1*–*3*). The cyclic GMP-AMP synthase (cGAS) is the primary cytosolic double-stranded DNA (dsDNA) sensor in mammalian cells (*4*–*7*). Upon binding to dsDNA, cGAS undergoes conformational changes that activate its ability to catalyze synthesis of a noncanonical 2’3’ cyclic-GMP-AMP dinucleotide (2’3’-cGAMP) that triggers type I interferon production through the endoplasmic reticulum membrane protein STING (also known as TMEM173, MPYS, MITA, and ERIS) (*8*–*13*). cGAS binds dsDNA in a sequence-independent manner (*14*–*20*). Abnormal activity of cGAS can lead to disease such as Aicardi-Goutières syndrome (*21*). cGAS activity is regulated by degradation or modification of the enzyme through ubiquitination, SUMOylation, phosphorylation, or glutamylation (*22*–*25*). A recent report suggests that cGAS activity is also regulated by the formation of phase-separated aggregates upon dsDNA engagement, which confine the activated cGAS to a particular location (*26*). Previous studies indicated that a variety of parameters, including the concentration and length of the dsDNA, influence sensitivity of cGAS-mediated detection of cytosolic DNA (*27*, *28*). However, cellular cGAS activity is not well explained by current structural and biophysical models (*26*, *28*).

Here we showed that cGAS activity is regulated through RNA-mediated phase separation. We found that cGAS forms phase-separated granules with RNAs as well as dsDNA. Aggregation with cytoplasmic RNAs, especially tRNAs, promotes enzyme activity of cGAS at low concentrations of dsDNA but inhibits cGAS enzyme activity when high concentrations of dsDNA are present. Thus, RNA plays a novel and important role in regulating cGAS activity.

### DNA- and RNA-induced phase separation of cGAS

We observed the formation of phase-separated granules soon after mixing the recombinant full-length human cGAS (FL-hcGAS) with a 45-bp double stranded interferon stimulatory DNA (ISD) (Figure S1A). Our results are consistent with the recently published work (*26*). In addition to dsDNA, previous studies showed that cGAS can bind RNAs, single-stranded DNAs (ssDNAs), and RNA-DNA hybrids (*29*–*31*). We also observed that the ssDNA without complementary regions can induce the formation of phase-separated granules when incubated with FL-hcGAS (Figure S1B). However, activation of FL-hcGAS was not observed when the ssDNAs had no complementary regions (Figure S1C). Short dsDNAs with one or more ssDNA arms induced strong phase separations of FL-hcGAS and activated the enzyme (Figure S2).

We next performed similar assays using total RNA extracted from HeLa cells. tRNAs are abundant in cytoplasm with an estimated concentration of approximately 1.2-1.8 mg/mL in mammalian cells and up to 20 mg/mL in yeast cells (*32*–*34*). Phase-separated granules were observed when FL-hcGAS was mixed with yeast tRNA and total RNA from HeLa cells over a wide RNA concentration range, although activation of FL-hcGAS was not detected (Figure 1). Phase separation of FL-hcGAS was observed after DNase treatment of the RNA preparation and in presence of a large amount of BSA (Figure S3); the later mimics the crowded environment in the cytoplasm. The binding of RNAs to FL-hcGAS was estimated using an electrophoretic mobility shift assay, and the results showed that RNA bound cGAS with a similar affinity as that of dsDNA (Figure S4), which is consistent with previous studies (*29*).

**Figure 1.**
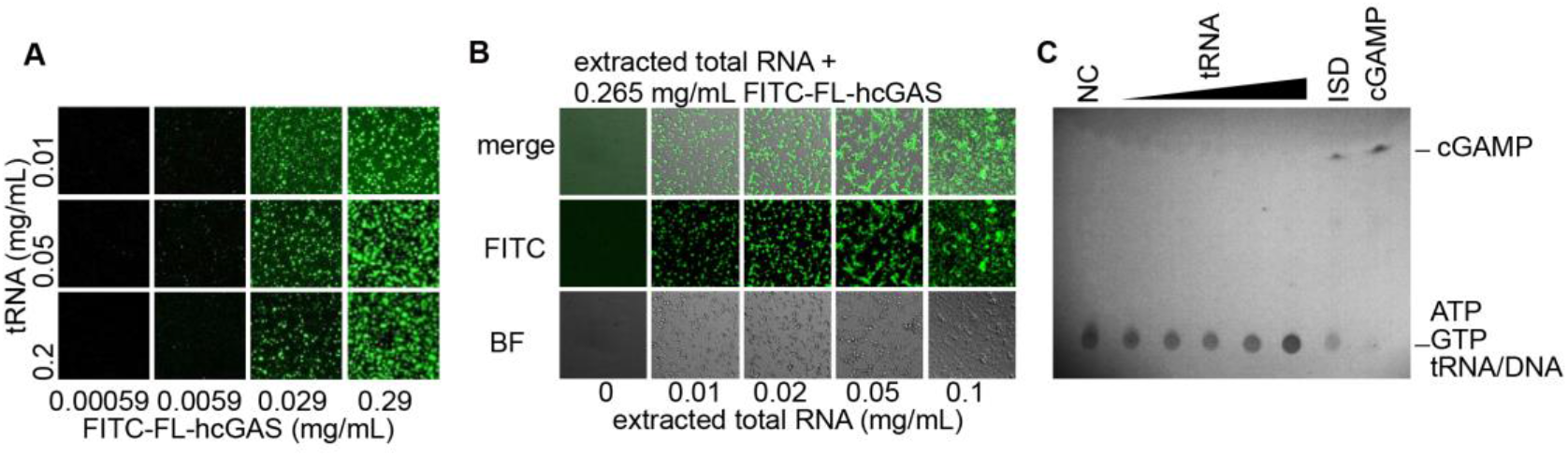
RNA mediates phase separation of cGAS. **A.** Fluorescent images of FITC-labeled full-length human cGAS (FITC-FL-hcGAS) incubated with yeast tRNA at the indicated concentrations. **B.** Fluorescent and bright field images of samples of FITC-FL-hcGAS (0.265 mg/mL) and total RNA from HeLa cells at the indicated concentrations. **C.** TLC analysis of cGAMP, which is indicative of the activation of cGAS. The FL-hcGAS concentrations was 0.265 mg/mL. Total RNA concentrations were 0.025, 0.05, 0.1, 0.25, and 0.85 mg/mL. NC indicates negative control which has only the FL-hcGAS. The dsDNA ISD was tested at 0.05 mg/mL as a positive control. The samples were prepared in 20 mM HEPES at pH 7.5 and 150 mM NaCl.

### RNA induces phase-separation of cGAS in cells

The concentration of RNAs, including tRNA and mRNA, in cytoplasm is much higher than is the concentration of DNA, probably even under abnormal conditions such as when cells are infected by viruses (*35*, *36*). To test the hypothesis that cGAS forms phase-separated granules with RNA in the absence of DNA, we over-expressed hcGAS as a C-terminal fusion with the fluorescent tag YFP in the HEK293T cells. The YFP-hcGAS gene was controlled under a doxycycline-inducible promoter and was integrated in the cell genome. Confocal microscopy revealed that a portion of the cells with the YFP signals had phase-separated granules in the cytoplasm (Figure 2A). Staining of the cells with Hoechst 33342 and pyronin Y, which as previously described stain DNA and RNA, respectively (*37*), showed that the YFP-hcGAS colocalized with cytoplasmic RNAs, especially in the granules of YFP-hcGAS (Figure 2A). In vitro assays showed that dsDNAs and RNAs in the cGAS-containing aggregates are differentially stained by using the combination of the two dyes (Figure S5).

**Figure 2.**
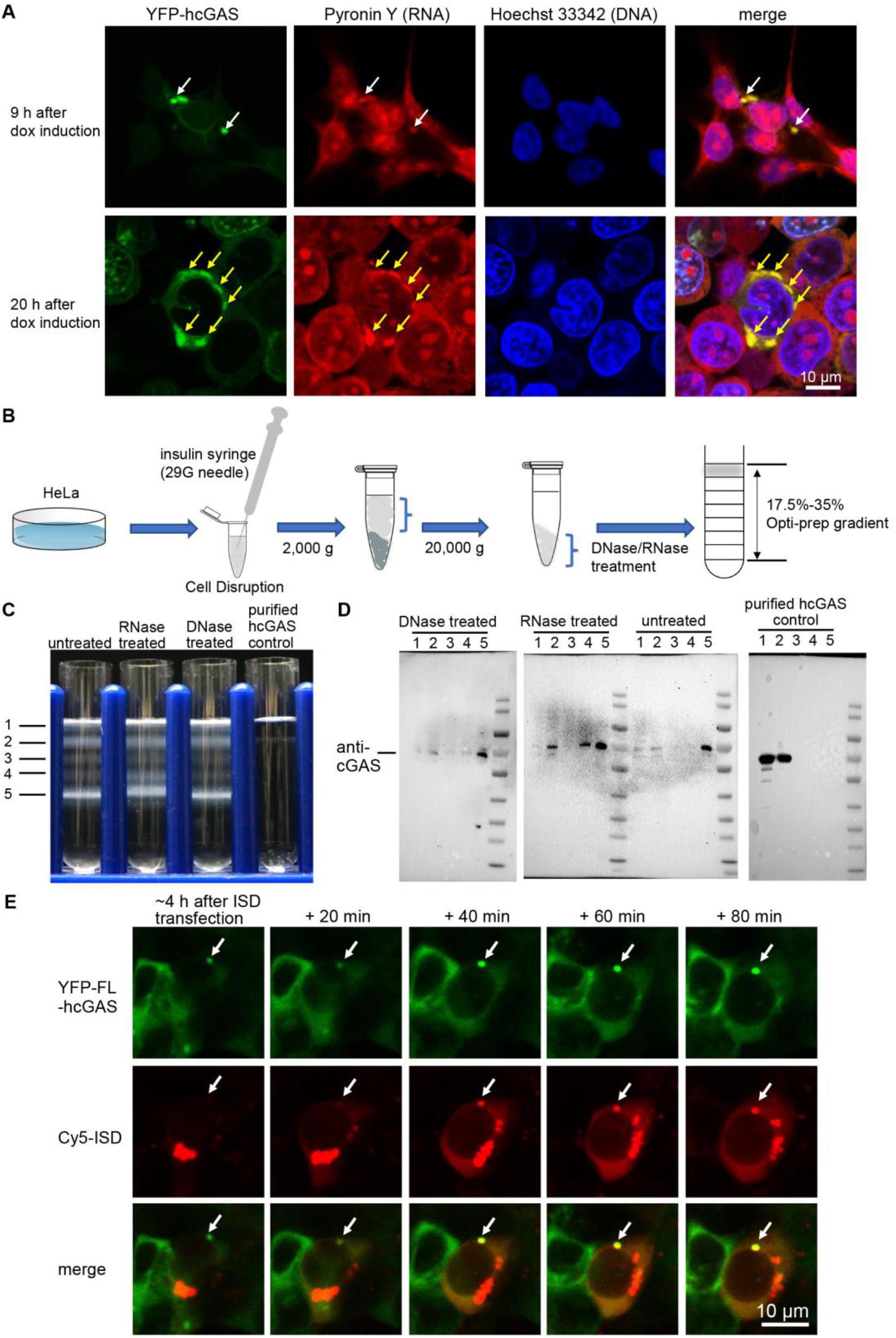
hcGAS associates with RNAs in cells. **A.** Hoechst 33342 and Pyronin Y staining of HEK293T cells showing colocalization of the YFP-cGAS granules and RNAs. Overexpression of the YFP-hcGAS was induced by doxycycline. **B.** A schematic diagram showing the extraction and fractionation procedure. To release the cytoplasm without disrupting the nuclear membrane, the cell membranes were disrupted in a hypotonic buffer by passing the cells through a 29G needle three times. The hypotonic buffer contained 10 mM HEPES, pH 7.5, 5 mM KCl, 3 mM MgCl_2_. **C.** Opti-prep gradient analysis of the HeLa cell cytoplasm extract. **D.** Western blotting analysis of samples of Opti-prep gradient bands with and without DNase and RNase treatment. **E.** Real time observation showing the eventual incorporation of the lipofectamine 2000 transfected Cy5-ISD into a preformed granule of YFP-hcGAS in a HEK293T cell.

The interactions between cGAS and RNAs in a cytoplasmic extract of Hela cells were analyzed using an Opti-prep gradient (Figure 2B). There were five major bands in the gradient after centrifugation (Figure 2C). hcGAS was detected in fractions from each of these bands by western blot with a cGAS-specific antibody. The endogenous cGAS proteins was located mainly in band 5. Band 5 was sensitive to RNase but not DNase (Figure 2C & 2D). Sequencing of the RNA in band 5 showed high numbers of reads for rRNAs and tRNAs (Figure S6), consistent with the abundance of different RNAs in cells (*33*). In the control gradient loaded with purified FL-hcGAS, the FL-hcGAS signals were detected by western blot mainly in fractions 1 and 2, near the top of the gradient (Figure 2C & 2D). These results indicate that endogenous hcGAS associates with RNAs in cells prior to sensing cytoplasmic dsDNAs, which are usually not present in the cytoplasm.

### dsDNA replaces RNA in preformed phase-separated granules

It was previously shown that molecules in the phase-separated granules are in dynamic equilibration with the molecules in solution (*38*). We observed that when Cy5-labeled ISD (Cy5-ISD) was transfected into HEK293T cells, the Cy5-ISD was eventually incorporated into the preformed granules of hcGAS-YFP (Figure 2E). As the electrophoretic mobility shift assays showed that the binding affinity of RNAs for hcGAS was comparable to that of dsDNA, in these cells where phase-separated granules exist prior to transfection, the transfected Cy5-ISD may replace RNA molecules in the granules.

To verify this, we performed in vitro assays with the fluorescein-5-thiosemicarbazide-labeled tRNA (FTSC-tRNA). Different amounts of ISD were added to solutions containing the preformed FTSC-tRNA-cGAS granules, and then the granules were separated from the solution by centrifugation and the signal due to FTSC-tRNA was measured in the supernatants (Figure 3A). As the concentration of the dsDNA was increased, the FTSC-tRNA signal increased in the supernatant until a plateau was reached (Figure 3B). This is indicative of a gradual substitution of the tRNAs by the dsDNAs until a dynamic equilibration was reached. Similarly, when granules were preformed from ISD and hcGAS, we observed a concentration dependent substitution of the dsDNAs by tRNAs (Figure 3C). Analyses by confocal microscopy also demonstrated that the nucleic acid component of the granules is in dynamic equilibrium (Figure 4).

**Figure 3.**
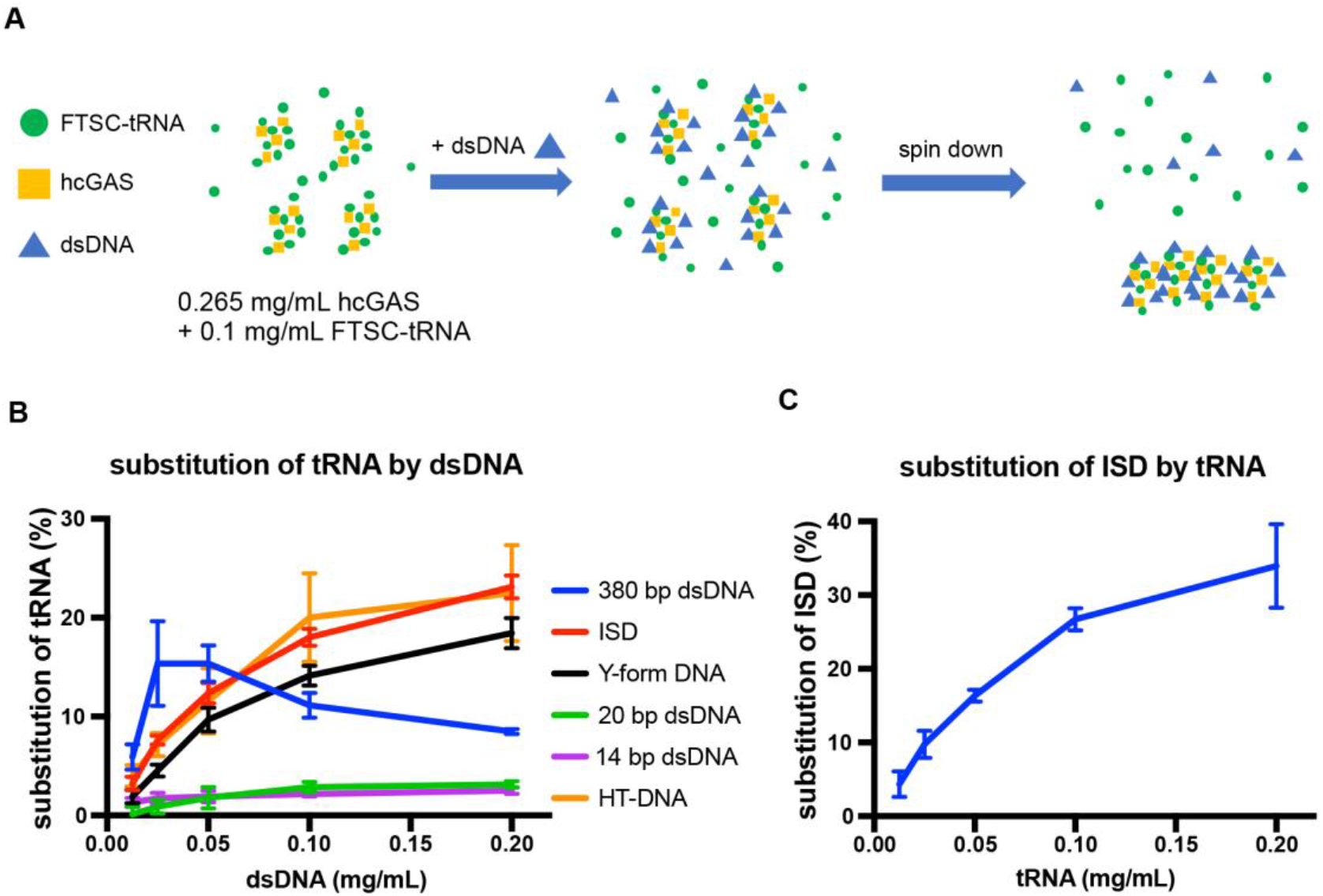
Substitution of the tRNA in the cGAS-tRNA phase separations by dsDNA. **A.** A schematic diagram showing the methods used to quantify tRNA/dsDNA released from the phase-separated granules. FTSC-tRNA and FL-hcGAS were pre-mixed. dsDNA was added, and the sample was centrifuged. The fluorescent signal of FTSC-tRNA in supernatant was measured. **B.** The percentage of FTSC-tRNA released in the supernatant as measured by fluorescence spectroscopy. The percentage of FTSC-tRNA released from the aggregates was calculated by using the following equation: P=(Fi-F0)*100/(F-F0). P: percentage of substituted FTSC-tRNA. Fi: fluorescence of FTSC-tRNA in the supernatant of each test. F0: fluorescence of FTSC-tRNA in the supernatant when DNA was not added. F: fluorescence of FTSC-tRNA without adding DNA and cGAS. The dsDNAs tested were ISD, 14 bp dsDNA, 20 bp dsDNA, 380 bp dsDNA, herring testis DNA (HT DNA) and Y-form DNA (a 14-bp dsDNA with unpaired GGG on each end). **C.** The percentage of Cy5-ISD released in the supernatant as measured by fluorescence spectroscopy. The yeast tRNA was used to trigger the release of the ISD from the phase separations of cGAS-ISD. The percentage of Cy5-ISD released from the aggregates was calculated by using the following equation: P=(Fi-F0)*100/(F-F0). P: percentage of substituted Cy5-ISD. Fi: fluorescence of Cy5-ISD in the supernatant of each test. F0: fluorescence of Cy5-ISD in the supernatant when tRNA was not added. F: fluorescence of Cy5-ISD without adding tRNA and cGAS. All experiments were performed three times.

**Figure 4.**
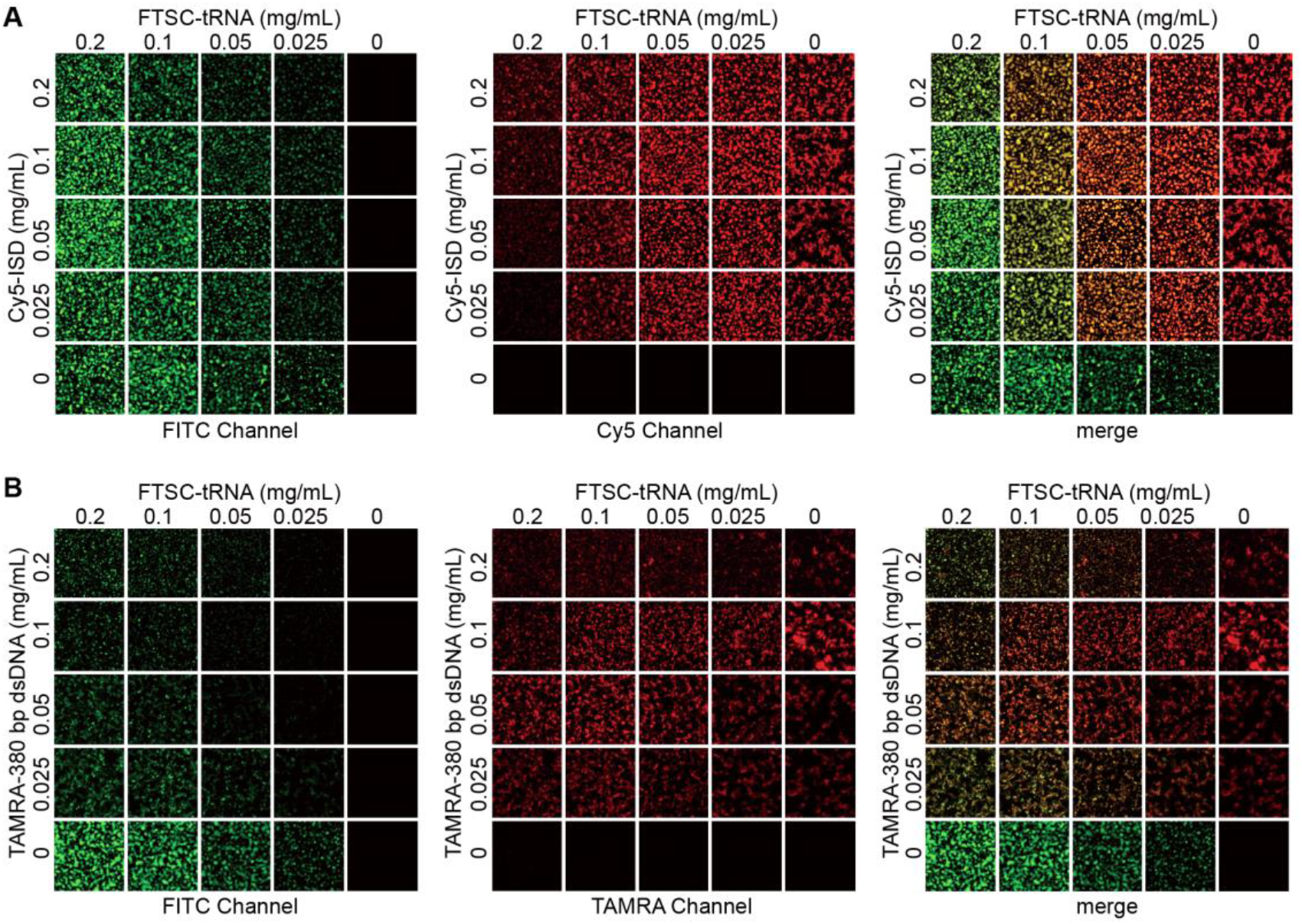
tRNA replacement by dsDNA in the cGAS-containing phase-separated granules observed by microscopy. **A.** Microscopy images of FTSC-tRNA in granules preformed with cGAS and indicated concentrations of FTSC-tRNA in the presence of indicated concentrations of ISD. **B.** Microscopy images of FTSC-tRNA in granules preformed with cGAS and indicated concentrations of FTSC-tRNA in the presence of indicated concentrations of a 380-bp dsDNA.

Long dsDNAs such as ISD, 380-bp dsDNA and herring testis DNA displaced the tRNA in the phase-separated granules even at a low concentration of 0.025 mg/mL (Figure 3B & Figure 4). In contrast, 14-bp and 20-bp dsDNAs, which bound to cGAS but did not induce the formation of aggregates (Figure S7), did not displace the tRNA (Figure 3B). A Y-form DNA, which is a 14-bp duplex with unpaired GGG at the termini, a structure previously shown to activate cGAS (*39*), induced the formation of the phase separations and also displaced tRNA from granules (Figure S7 and Figure 3B).

### tRNA-induced phase separation regulates cGAS activity

Since hcGAS is not activated by binding to RNA, we reasoned that the competitive binding of RNAs to hcGAS should have a negative impact on the dsDNA-dependent activation of the enzyme. We measured the hcGAS activity with or without tRNA and showed that cGAS activity was significantly inhibited by tRNA when DNA concentration exceeded 0.05 mg/mL (Figure 5A). tRNA had little or no effect on cGAS activity when the DNA concentration was less than 0.01 mg/mL, however (Figure 5A). At a high dsDNA concentration (0.0544 mg/mL), tRNA inhibited production of 2’3’-cGAMP catalyzed by hcGAS (Figure 5B), whereas at a low dsDNA concentration (0.0068 mg/mL) enzyme activity was stimulated (Figure 5C). Robust phase separation was observed at high DNA concentration (Figure 5D) but not at low DNA concentration (Figure 5E). The phase separation-related turbidity of the solution did not change significantly as a function of tRNA concentration with a high concentration of dsDNA (Figure 5F). However, the fluorescent signal due to FAM-labeled dsDNA was reduced within the granules upon addition of higher concentrations of tRNA (Figure 5D & 5H). The phase separation related turbidity of the solution increased significantly as a function of tRNA concentration when the tRNA was added with a low concentration of dsDNA (Figure 5G). The fluorescent signal in the phase-separated granules remains constant. Thus, tRNA promotes the formation of phase separation when the dsDNA concentration is not high to induce aggregation (Figure 5E & 5I). This tRNA mediated formation of phase separation promotes the activation of cGAS with even only a few dsDNA molecules (Figure 5C).

**Figure 5.**
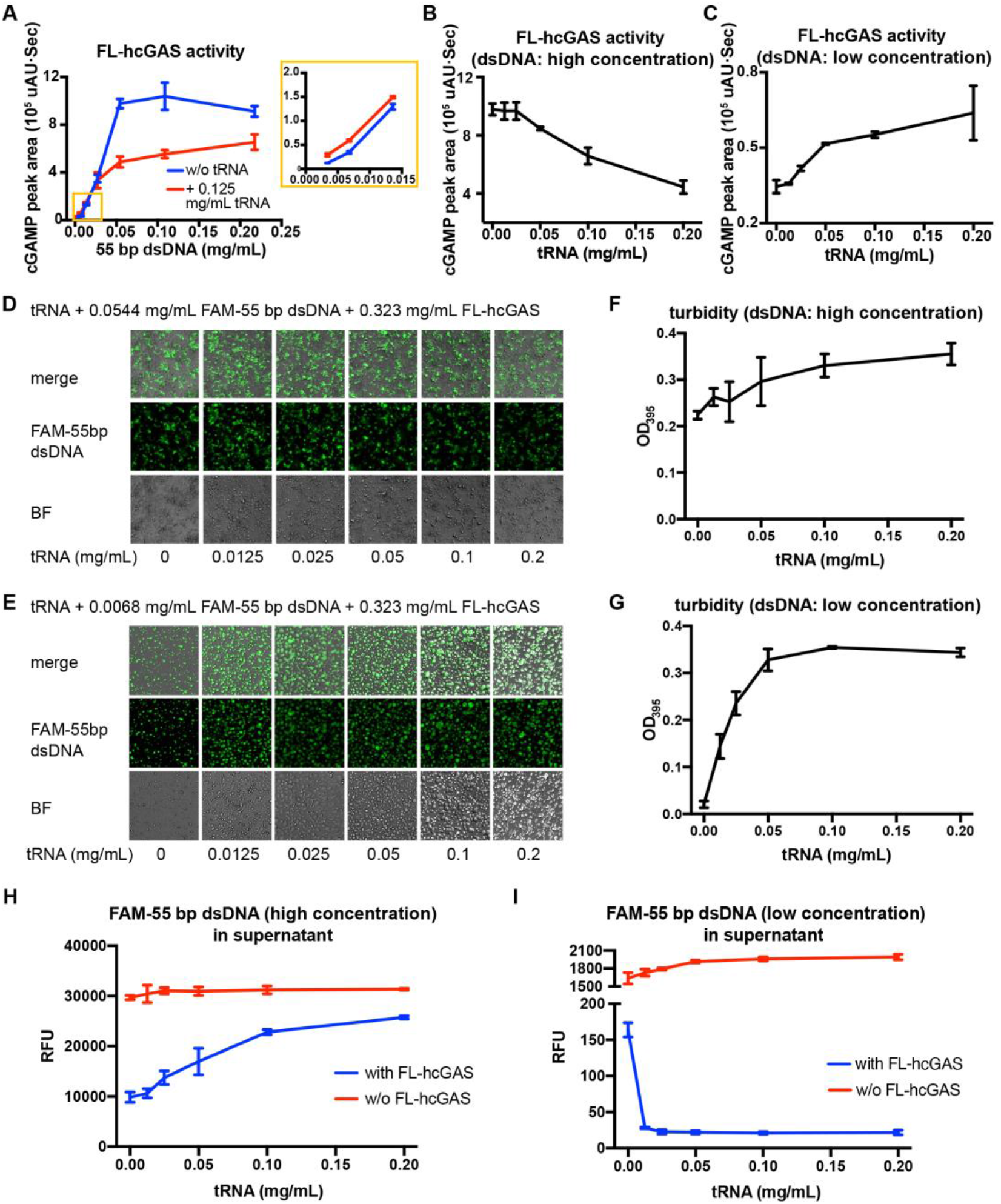
tRNA regulates cGAS activity. **A.** cGAS activity (as measured by cGAMP peak area) with or without 0.125 mg/mL tRNA at indicated concentrations of a 55-bp dsDNA. The inset is an expanded view of the region boxed in yellow. **B.** cGAS activity in presence of 0.0544 mg/mL of 55-bp dsDNA as a function of tRNA concentration. **C.** cGAS activity in presence of 0.0068 mg/mL of 55-bp dsDNA as a function of tRNA concentration. **D.** Fluorescent and bright field photographs of phase-separated granules of FL-hcGAS and 0.0544 mg/mL of FAM-labeled 55-bp dsDNA at the indicated tRNA concentrations. **E.** Fluorescent and bright field photographs of phase-separated granules of FL-hcGAS and 0.0068 mg/mL of FAM-labeled 55-bp dsDNA at the indicated tRNA concentrations. **F.** Turbidities of samples of FL-hcGAS and 0.0544 mg/mL of FAM-labeled 55-bp dsDNA at the indicated tRNA concentrations measured at 395 nm. **G.** Turbidities of samples of FL-hcGAS and 0.0068 mg/mL of FAM-labeled 55-bp dsDNA at the indicated tRNA concentrations measured at 395 nm. **H.** Fluorescent signals of the FAM-labeled 55-bp dsDNA in the supernatants of the samples of FL-hcGAS and 0.0544 mg/mL of FAM-labeled 55-bp dsDNA at the indicated tRNA concentrations. **I.** Fluorescent signals of the FAM-labeled 55-bp dsDNA in the supernatants of the samples of FL-hcGAS and 0.0068 mg/mL of FAM-labeled 55-bp dsDNA at the indicated tRNA concentrations. Each condition was evaluated in triplicate.

## Discussion

By combining biochemical and cellular assays, we have unequivocally established that cGAS forms phase-separated granules with RNAs in cytoplasm of human cells and that these RNA-containing granules regulate the sensitivity of cGAS activity to cytosolic dsDNA. At a low dsDNA concentration that is not enough to induce the formation of phase separation, cytoplasmic RNAs, especially tRNAs, form aggregates with cGAS that provide platforms for dsDNA-mediated cGAS activation. When the cytoplasmic concentration of dsDNA is high enough to induce phase separation and activate cGAS, tRNAs compete with dsDNA to bind cGAS and inhibit cGAS activity, presumably to limit of over activation of the enzyme. Given the high concentration of the RNAs in cytoplasm, the RNAs are likely the dominant regulators of cGAS activity. Our observation offers a reasonable mechanism by which cGAS sensitively detects cytosolic dsDNA but is modulated to ensure an appropriate immune response to cytosolic dsDNA. Our finding that short dsDNAs, such as 14- and 20-bp duplexes do not efficiently displace RNAs from phase-separated granules provides an explanation of why short dsDNAs do not activate cGAS in cells, although they can active cGAS in vitro (*39*, *40*). The abundances of tRNAs and mRNAs are altered under stress conditions (*41*), suggesting that RNAs could have complex functions in regulation of cGAS-mediated innate immune responses.

## Materials and methods

### Protein expression and purification

The coding sequence of *hcGAS* was optimized for *E. coli* expression using the GeneOptimizer algorithm (Thermo). The synthetic gene (Qinglan) was cloned into a modified pETDuet at a site designed to fuse a 10x His tag and a sumo tag at the N-terminus of the protein. The pETDuet-hcGAS plasmid was transformed into *E. coli* BL21 Star (DE3) competent cells. The transformed cells were cultured in six 800-mL aliquots of LB at 37 °*C* until the absorbance at 600 nm reached ~ 0.6. The cells and medium were cooled to 16 °C, and 1 mM IPTG was added to induce protein expression. The cells were harvested 16 h after the induction and were resuspended in 100 mL PBS buffer at pH 7.0 with 300 mM NaCl. The resuspended cells were homogenized and the cell lysate was centrifugated at 20,000 g for 15 min. The supernatant was applied to 4-mL Talon Metal Affinity Resin (Clontech, cat# 635503). After washing with 30 mL wash buffer containing 20 mM HEPES at pH 7.5, 150 mM NaCl, and 10 mM imidazole, the resin was resuspended in 11 mL of buffer containing 20 mM HEPES, pH 7.5, 150 mM NaCl, and 0.1 mg/mL ULP1 and incubated at 4 °*C* for 12 h to remove the sumo tag. The hcGAS released from the resin was purified over a heparin column (GE Healthcare) to remove any dsDNA contamination and was further purified over a Superdex 75 size-exclusion column (GE Healthcare) using a running buffer containing 20 mM HEPES, pH 7.5, and 150 mM NaCl.

### Fluorescence labeling of the tRNA, dsDNA, ssDNA, cGAS

Yeast tRNAs (Solarbio, cat# T8630) were labeled with FTSC as described by Qiu et al. (*42*). In brief, a solution containing 20 μL tRNA (10 mg/mL) was mixed with 200 μL of 0.25 M sodium acetate. The mixture was diluted with ddH_2_O to a final volume of 700 μL, and 50 μL of 1 mM NaIO_4_ was added to oxidize the tRNAs. After incubation for 90 min in the dark at room temperature, the oxidization reaction was stopped by adding 40 μL of 2.5 mM Na_2_SO_3_ to the solution followed by incubation for 15 min at room temperature. Labeling of the tRNAs was performed by adding 60 μL of 2.5 mM FTSC in DMF, and the solution was incubated for 3 h in the dark at room temperature. Excess FTSC was removed using a 10-kDa cutoff Ultra Centrifugal Filter (Millipore) using 20 mM HEPES, pH 7.5, and 150 mM NaCl as the buffer.

ISD and the 55-bp dsDNA were labeled by primer extension with the forward primer conjugated to FAM or Cy5 on the 5’ terminus (Sangon). The ISD sequence is 5’-TACAGATCTACTAGTGATCTATGACTGATCTGTACATGATCTACA-3’. The 55-bp dsDNA sequence is 5’-TCGATACAGATCTACTAGTGATCTATGACTGATCTGTACATGATCTACAAT CACT-3’. The 380-bp dsDNA was amplified from the SARS-CoV genome using a forward primer with a 5’ TAMRA label (Sangon). The primer sequences are forward 5’-TAATACGACTCACTATAGGGATGTCTGATAATGG-3’ and reverse 5’-AGCTTCTGGGCCAGTTCCTAG-3’.

The cGAS protein was labeled with FITC (Thermo, cat# 46424). The protein in 20 mM HEPES, pH 7.5, and 150 mM NaCl at a concentration of 1-5 mg/mL and FITC were mixed at a molar ratio of 1:1 to 1:1.5. After incubation for 5 min in the dark at room temperature, 2 mM Tris-HCl, pH 8.0 was added to the solution to stop the reaction. Protein aggregates were removed by centrifugation at 20,000 g for 5 min, and excess FITC was removed using a 30-kDa cutoff Ultra Centrifugal Filter (Millipore) with the 20 mM HEPES, pH 7.5, and 150 mM NaCl as buffer.

### Observation of phase-separated aggregates

For the phase separations induced by the total RNA from HeLa cells or yeast tRNA, RNAs and FITC-labeled cGAS were mixed at the indicated concentrations and were transferred into a 384-well plate. The samples were prepared in 20 mM HEPES at pH 7.5 and 150 mM NaCl. Total RNA was extracted from HeLa cells with an RNA purification kit (Thermo, cat# K0731). Phase separations of the RNAs and FITC-labeled cGAS were observed by using laser scanning confocal microscopy with excitation set at 488 nm and emission filter set at 499-641 nm.

For substitution of FTSC-tRNA in cGAS-associated phase-separated granules by dsDNAs, FTSC-tRNA and cGAS were mixed first and then dsDNAs (Cy5-ISD or TAMRA-380 bp dsDNA) were added at the indicated concentrations. The samples were prepared in 20 mM HEPES at pH 7.5 and 150 mM NaCl. The samples were transferred into a 384-well plate. Observation was achieved by using laser scanning confocal microscopy. Excitation wavelengths and emission filters were set as follows: FTSC excitation: 488 nm; FTSC emission filter: 491-535 nm; Cy5 excitation: 633 nm; Cy5 emission filter: 638-759 nm; TAMRA excitation: 543 nm; TAMRA emission filter: 558-682 nm.

To observe the influence of tRNA on the phase separation of cGAS and dsDNA, FAM labeled 55-bp dsDNA, tRNA and cGAS were mixed at the indicated concentrations. The samples were prepared in 25 mM Tris-HCl at pH 8.0, 20 mM NaCl, 5 mM MgCl2, 1 mM ATP and 1 mM GTP. Phase separation was observed by using laser scanning confocal microscopy with excitation set at 488 nm and emission filter set at 499-641 nm.

### Stable cell line generation

To generate the cell line that stably expresses doxycycline-inducible YFP-hcGAS, the sequence encoding YFP-hcGAS was cloned into the lentiviral vector pLVX-TetOne-Puro. A mixture of 2 μg pLVX-TetOne-Puro-YFP-hcGAS, 1 μg pMD2.G, and 1 μg psPAX2 was transfected into the HEK293T cells using Lipofectamine 2000 (Thermo, cat# 11668019) when cells were at ~70% confluence. The cells were cultured in a 6-well plate with the DMEM medium (Thermo) and 10% FBS (Gibco, cat# 10091148) to generate the lentivirus. The medium containing lentiviruses was collected 60 h after the transfection, and the dead cell debris were removed by centrifugation. Polybrene was added to the solution at a final concentration of 0.8 μg/mL to enhance the lentiviral infection. The lentivirus was added to HEK293T cells cultured in a 6-well plate when cells were at 50-60% confluence. Virus was replaced 12 h after infection with fresh DMEM medium containing 10% FBS. The cells were cultured for another 12 h, and then puromycin was added to the medium at a final concentration of 2.5 μg/mL. After passaging the cells three times in medium containing 2.5 μg/mL puromycin, the cells were maintained in the medium containing 2.5 μg/mL puromycin.

### Cell imaging

For RNA/DNA staining and imaging, YFP-hcGAS HEK293T cells were seeded on a coverslip. After 24 h, expression of YFP-hcGAS was induced by addition 0.1 μg/mL doxycycline, and cells were allowed to grow for 9-20 h. Cells were fixed with 4% polyformaldehyde in PBS and were washed with PBS after 15 min. The cells were permeabilized with 0.5% Triton X-100 in PBS for 15 min, and then the cells were washed three times with PBS. PBS containing 2 μg/mL Hoechst 33342 and 4 μg/mL pyronin Y (Amresco, cat# 0207) was added to the cells to stain DNA and RNA, respectively. After 15 min, coverslips were mounted on glass slides with Thermo Prolong Glass Antifade Mountant. After 2 h or longer, the slides were observed by confocal microscopy (Zeiss LSM 880). For each dye, excitation lasers and emission filters were as follows: Hoechst 33342 excitation, 405 nm; Hoechst 33342 emission, 410-489 nm; YFP excitation, 514 nm; YFP emission, 525-588 nm; pyronin Y excitation, 561 nm; and pyronin Y emission, 625-758 nm. Phase-separated granules of cGAS and the 55-bp dsDNA, total RNA, or tRNA were observed as controls (Figure S5).

For imaging of YFP-hcGAS HEK293T cells transfected with Cy5-ISD, cells were cultured in a four-chamber glass bottom dish (Cellvis, cat# D35C4-20-1.5-N). After expression was induced by treatment with 0.1 μg/mL doxycycline for 6-9 h, 0.5 μg of Cy5-ISD was transfected into the cells in one chamber using Lipofectamine 2000. Cells were observed using a confocal microscope (Zeiss LSM 710).

### tRNA displacement assays

FTSC-labeled tRNA and cGAS were mixed to allow tRNA-cGAS aggregates to form, and then DNA was added. After a short incubation of 5 min at room temperature, cGAS-nucleic acid aggregates were spun down at 20,000 g for 5 min. The fluorescence signal of the free FTSC-tRNA in the supernatant was measured using a NanoDrop 3300 (Thermo Scientific). Release of the FAM-labeled ISD from the aggregates was measured in a similar way. The concentrations of the FTSC-tRNA and of the FAM-dsDNA had linear relationships with fluorescent signals in the range of 0.0125 to 0.2 mg/mL (Figure S8). The percentage of FTSC-tRNA displaced in the aggregates was calculated by using the following equation: P=(Fi-F0)*100/(F-F0). P: percentage of the substituted FTSC-tRNA. Fi: fluorescence signal of FTSC-tRNA in the supernatant of each test. F0: fluorescence signal of FTSC-tRNA in the supernatant when DNA was not added. F: fluorescence signal of FTSC-tRNA without adding DNA and cGAS. The percentage of Cy5-ISD displaced in the aggregates was calculated by using a similar equation.

### Cytoplasmic cGAS isolation and density gradient centrifugation

The isolation of cytoplasmic cGAS from HeLa cells was previously described (*26*). The cytoplasm was extracted by hypotonic treatment (10 mM HEPES, pH 7.5, 5 mM KCl, 3 mM MgCl_2_) and homogenization through a 26G needle. The homogenized cells were centrifuged at 2,000 g to remove cell debris and nuclei. The supernatant (S2) was collected and centrifuged at 20,000 g to obtain the supernatant fraction S20 and the pellet fraction (P20). P20 was resuspended in an isotonic buffer (20 mM HEPES, pH 7.5, 250 mM sucrose, 25 mM KCl, 5 mM MgCl_2_) containing 17.5 % Opti-prep (Sigma, cat# D1556). The resuspended P20 (500 μL) was further fractionated over an Opti-prep density gradient after treatment with RNase (20 μL of 10 mg/mL RNase A, 5 μL of 10 U/μL RNase I), DNase (25 μL of 1 U/μL DNase I), or buffer for 30 min at 37 °C. After further fractionation over an Opti-prep density gradient (isotonic buffer containing 35%, 32.5%, 30%, 27.5%, 25%, 22.5%, or 20% Opti-prep, from bottom to top, 500 μL per layer, with resuspended P20 on the top of the gradient).

### In vitro analysis of cGAS activity

In vitro enzymatic activities of cGAS were measured by monitoring the formation of the product 2’3’-cGAMP by using a HPLC system (SHIMADZU, LC-10A) equipped with LC-10AT pumps and an SPD-10AV ultraviolet detector. The strands of the 55-bp dsDNA were chemically synthesized: 5’-TCGATACAGATCTACTAGTGATCTATGACTGATCTGTACATGATCTACAAT CACT-3’ and 5’-AGTGATTGTAGATCATGTACAGATCAGTCATAGATCACTAGTAGATCTGTA TCGA-3’. The reaction mixture contained 5.5 μM (0.323 mg/mL) FL-hcGAS, 25 mM Tris-HCl, pH 8.0, 20 mM NaCl, 5 mM MgCl_2_, 1 mM ATP, 1 mM GTP, the 55-bp dsDNA at various concentrations (0.0034 – 0.2166 mg/mL) with or without 5 μM (0.125 mg/mL) tRNA. Reactions were also performed by varying tRNA concentrations (0.5-8 μM, 0.005-0.200 mg/mL) at 0.0068 or 0.0544 mg/mL 55-bp dsDNA with all other components the same as described above. Each reaction in 40 μL total volume was incubated at 25 °*C* for 5 min, and terminated by heating at 85 °C for 15 min. After centrifuging at 16,000 g for 10 min, the supernatant was analyzed using an YMC-pack pro-C18 reverse phase column (4.6 x 250 mm, 5 μm). Analytes were monitored using 254 nm light. Buffer A contained 5 mM ammonium acetate (pH 5.0) in water, and phase B was 100% acetonitrile. The samples were eluted using a linear gradient from 2% B to 15% B at a flow rate of 1 mL/min over 15 min. The product 2’3’-cGAMP was quantified by measuring the peak area (uAU·sec) of 2’3’-cGAMP visualized with 254 nm light. Each experiment was performed in triplicate.

## Supplemental information

**Supplemental Figure 1.**
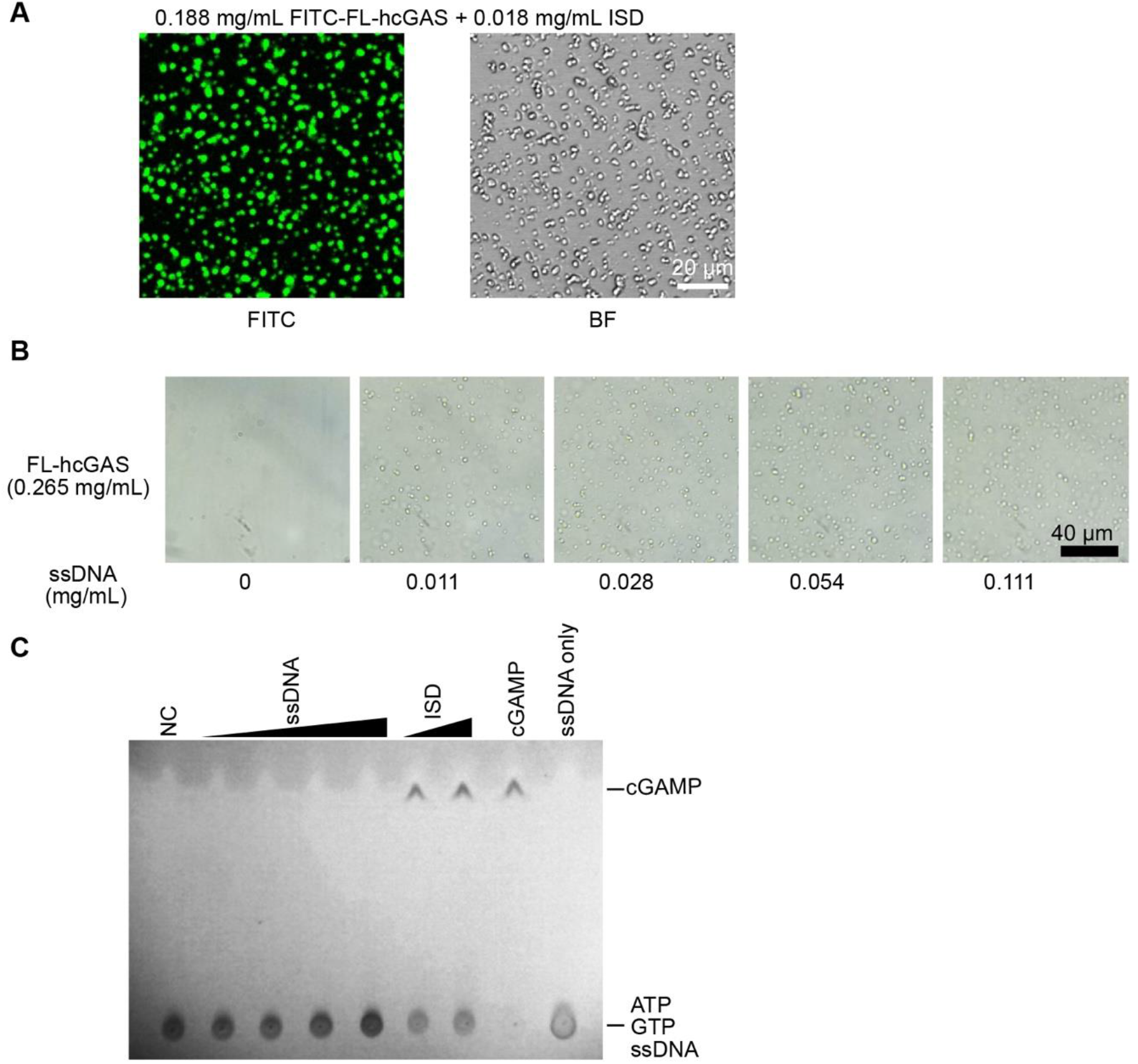
ssDNA mediates phase separation of FL-hcGAS. **A.** Fluorescent and bright field photographs showing phase-separated granules of FITC-labeled FL-hcGAS induced by the dsDNA referred to as ISD (for interferon stimulatory DNA), with the sequence 5’-TACAGATCTACTAGTGATCTATGACTGATCTGTACATGATCTACA-3’. The FITC-labelled FL-cGAS concentration was 0.188 mg/mL and the ISD concentration was 0.018 mg/mL. **B.** Bright field photographs showing samples of FL-hcGAS (0.265 mg/mL) and a 48 nt ssDNA at the indicated concentrations. The 48 nt ssDNA has the sequence 5’-AGAGAGAGAGAGAGAGAGAGAGAGAGAGAGAGAGAGAGAGAGAGAGAG-3’. **C.** TLC analysis for the presence of cGAMP, which is indicative of cGAS activation. The FL-hcGAS concentrations was 0.265 mg/mL. ssDNA concentrations were 0.011, 0.028, 0.054 and 0.111 mg/mL. NC indicates negative control which has only the FL-hcGAS. The dsDNA ISD was tested at 0.055 and 0.111 mg/mL as positive controls. Samples were prepared in 20 mM HEPES at pH 7.5 and 150 mM NaCl.

**Supplemental Figure 2.**
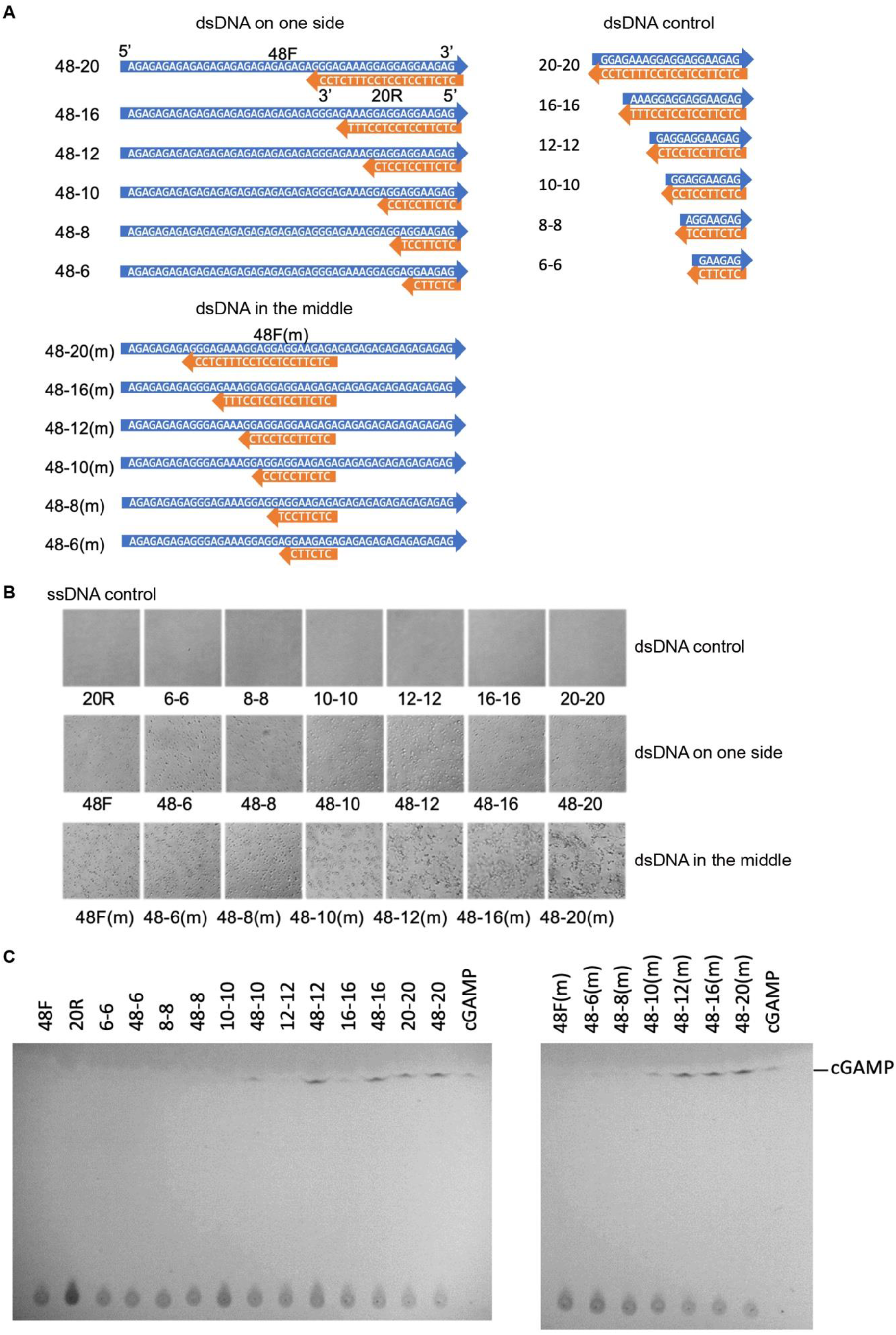
Short dsDNAs with ssDNA arms promote phase separation and activation of cGAS. **A.** Sequences of short dsDNAs with ssDNA arms and dsDNA controls. **B.** Photographs of samples of FL-hcGAS (0.265 mg/mL, 4.4 μM) with indicated DNAs (2 μM) taken with a differential interference contrast microscope. **C.** TLC analysis for cGAMP, which is indicative of the activation of cGAS, in the presence of short dsDNAs with ssDNA arms. The FL-hcGAS and DNA concentrations were 4.4 μM and 2 μM, respectively. Samples were prepared in 20 mM HEPES at pH 7.5 and 150 mM NaCl.

**Supplemental Figure 3.**
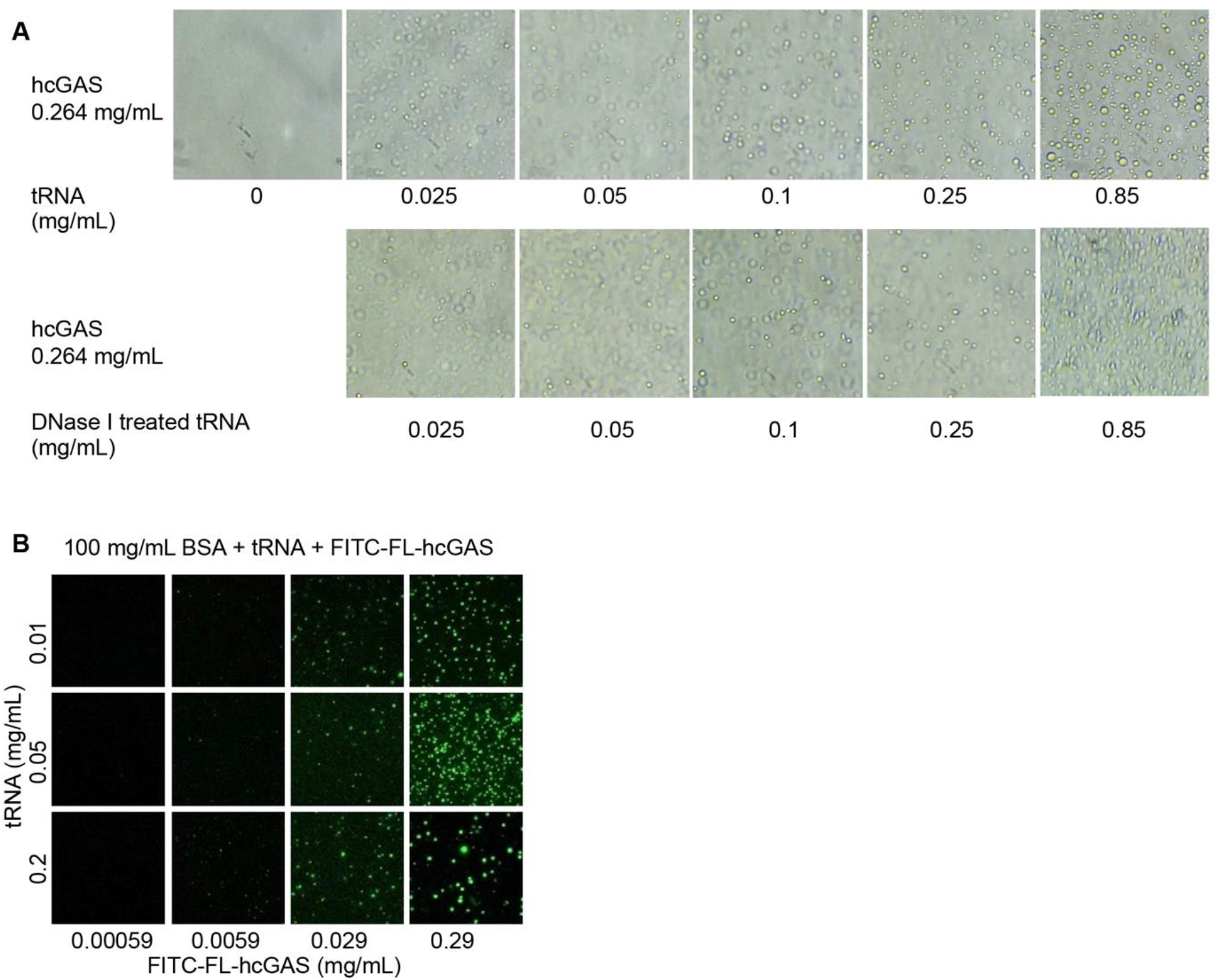
RNA mediates phase separation of cGAS. **A.** Upper: Bright field photographs of samples of FL-hcGAS (0.264 mg/mL) and total RNA from HeLa cells at the indicated concentrations. Lower: Bright field photographs of samples of FL-hcGAS (0.264 mg/mL) and DNase I-treated total RNA at the indicated concentrations. **B.** Fluorescent images of FITC-labeled FL-hcGAS and total RNA from HeLa cells in presence of 100 mg/mL BSA; concentrations of FL-hcGAS and total RNA are indicated. Samples were prepared in 20 mM HEPES, pH 7.5, 150 mM NaCl.

**Supplemental Figure 4.**
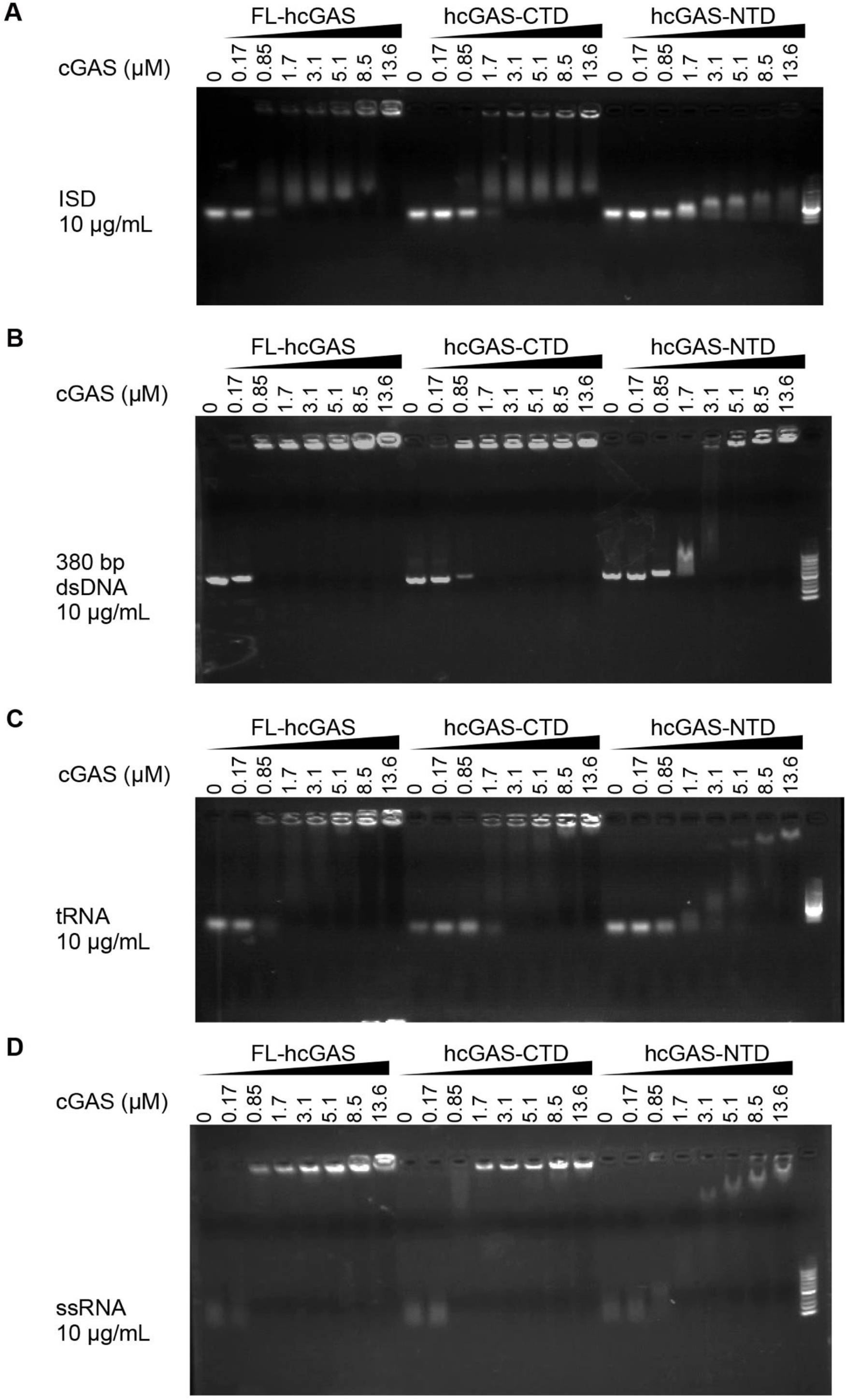
RNA and DNA bind cGAS with similar affinity. Electrophoretic mobility shift analysis of ISD, 380-bp dsDNA, yeast tRNA and ssRNA in presence of FL-hcGAS. The nucleic acid concentration was 10 ng/μL. The cGAS concentrations were 0, 0.17, 0.85, 1.7, 3.4, 5.1, 8.5, and 13.6 μM.

**Supplemental Figure 5.**
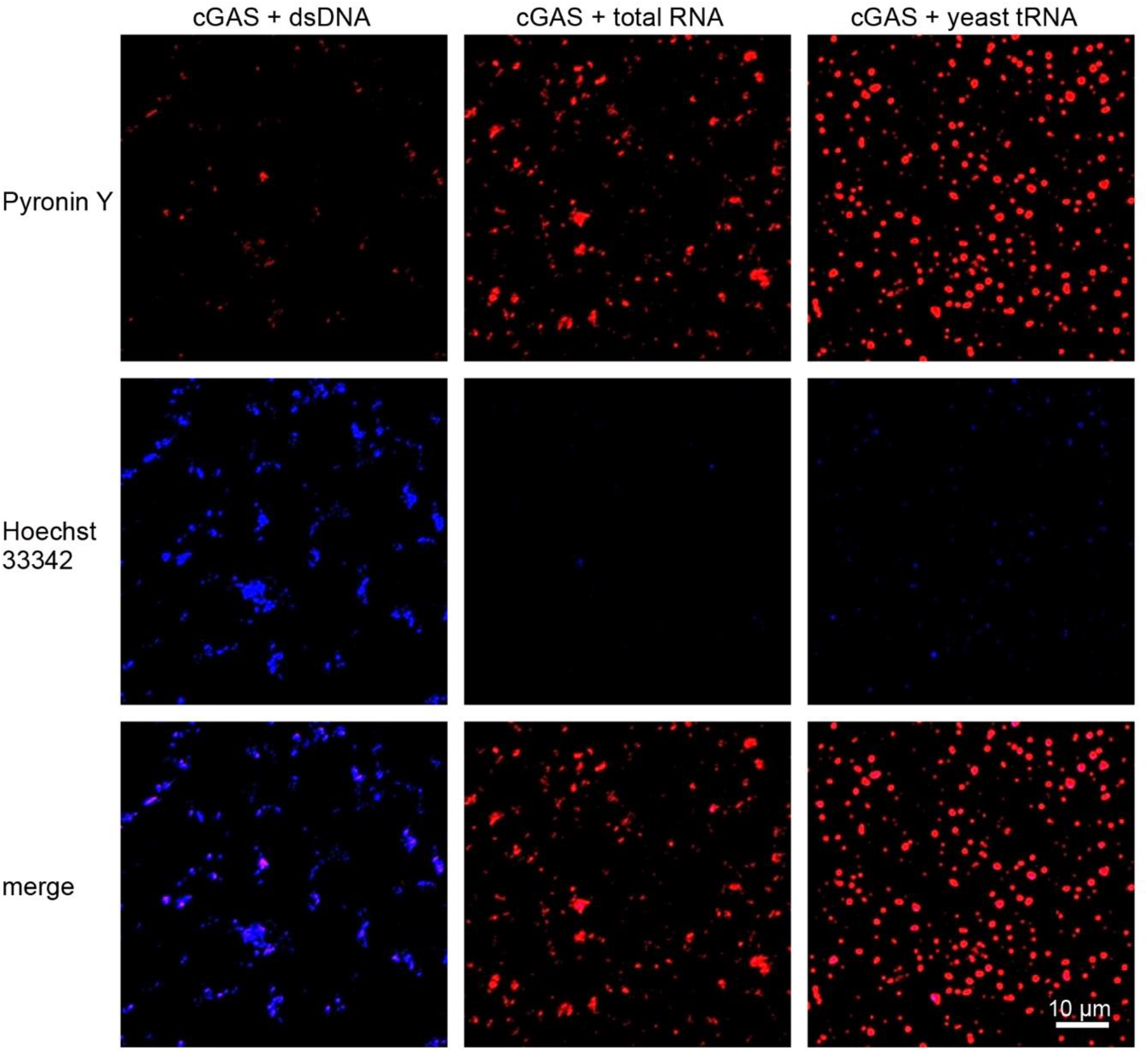
Hoechst 33342 and pyronin Y differentially stain dsDNA and RNA in the cGAS involved phase separations. Confocal microscopy images of FL-hcGAS incubated with 55-bp dsDNA, total RNA from HeLa cells, or yeast tRNA and stained with 2 μg/mL Hoechst 33342 and 4 μg/mL pyronin Y. The FL-hcGAS concentrations was 0.2 mg/mL. 55-bp dsDNA, total RNA and yeast tRNA concentrations were both 0.1 mg/mL.

**Supplemental Figure 6.**
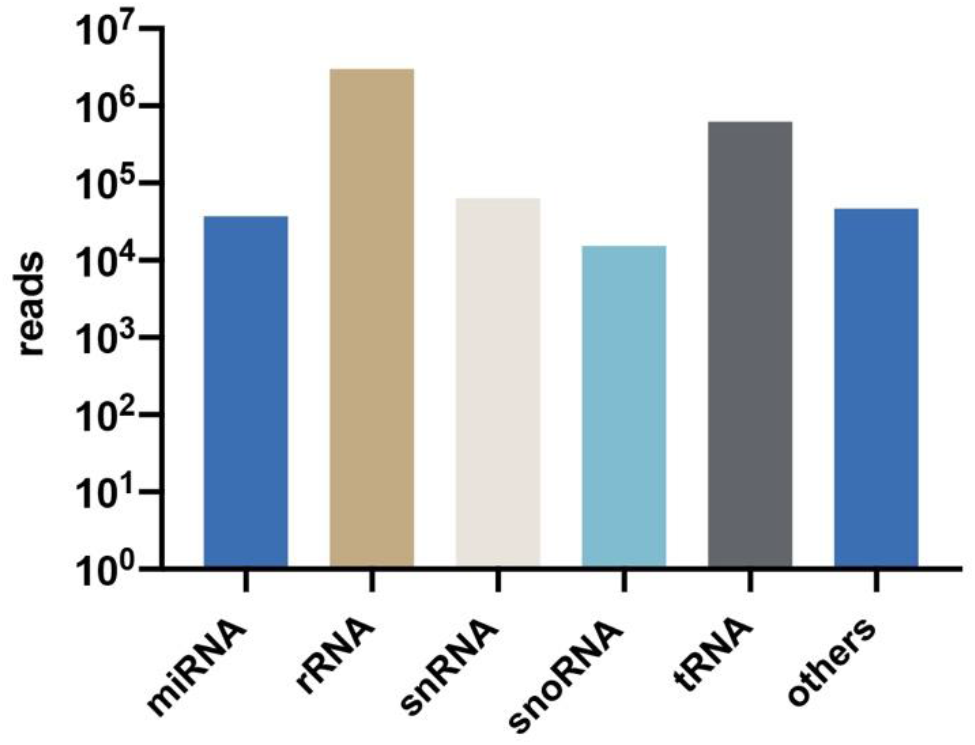
Sequencing of RNA in the cGAS-containing fraction from cytoplasm. Two libraries were generated from band 5 from the Opti-prep gradient: one for non-coding RNA and one for mRNA sequencing. The histograms show the numbers of non-coding RNA sequencing reads. mRNA was also sequenced and the results showed coverage of most of the actively transcript genes.

**Supplemental Figure 7.**
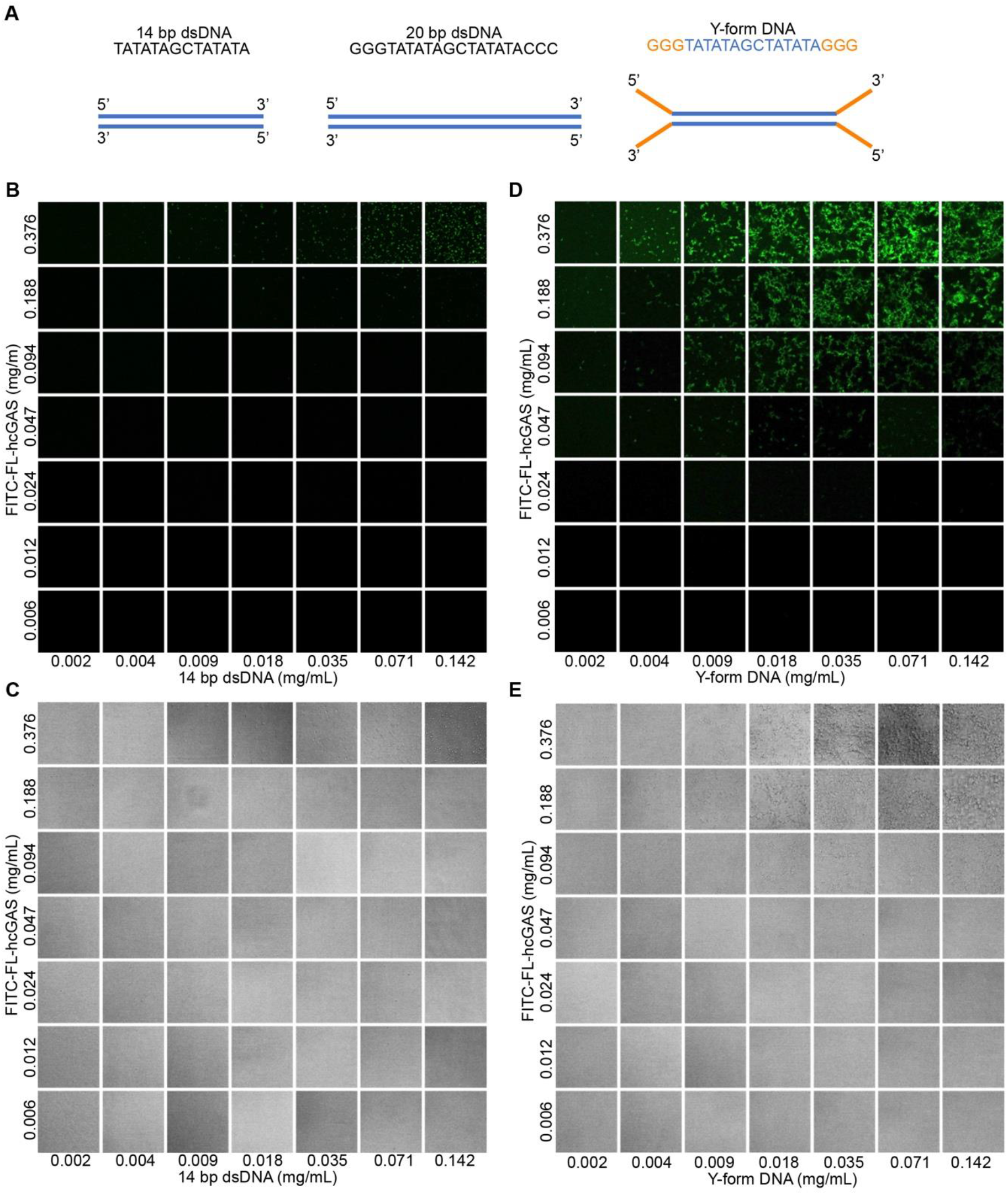
Y-form DNA and 14-bp dsDNA mediate phase separation of FL-hcGAS. **A.** Sequences and structures of the, 14-bp dsDNA, 20-bp dsDNA and Y-form DNA. **B.** Fluorescent images of samples of FITC-FL-hcGAS and the 14-bp dsDNA at indicated concentrations. **C.** Corresponding bright field photographs of samples of FITC-FL-hcGAS and the 14-bp dsDNA at indicated concentrations. **D.** Fluorescent images of samples of FITC-FL-hcGAS and the Y-form DNA at indicated concentrations. **E.** Corresponding bright field photographs of samples of FITC-FL-hcGAS and the Y-form DNA.

**Supplemental Figure 8.**
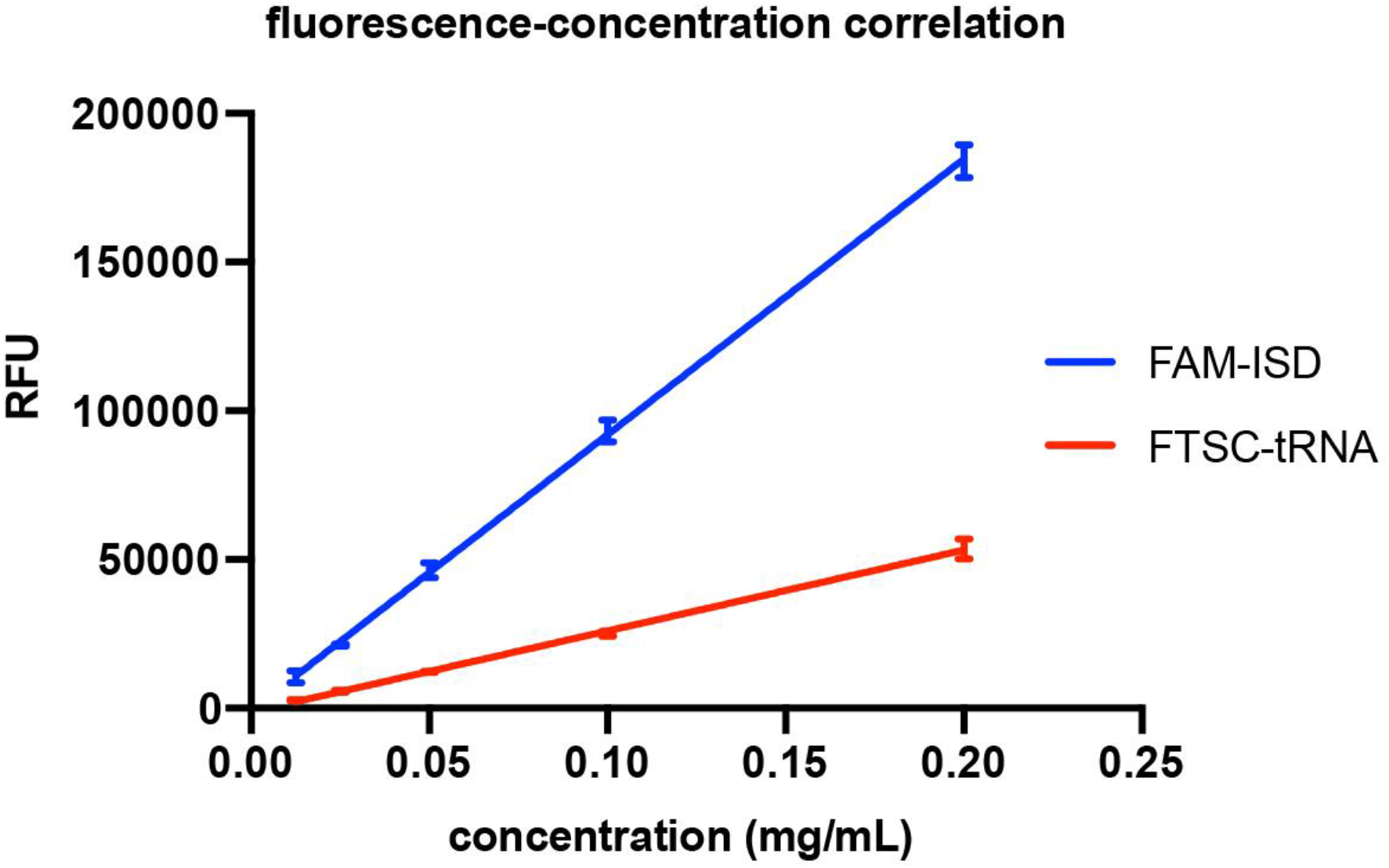
Correlation of fluorescence signal (RFU) with concentration for fluorophore labeled ISD and tRNA.

## Author Information

The authors declare no competing financial interests. Correspondence and requests for materials should be addressed to D.Y.Z.; Y. X..

## Author Contributions

X.Y. designed the research; S.L.C., D.Z, M.R., Y. L., and X.Y. performed the experiments; S.L.C., D.Z., and X.Y. analyzed data and wrote the paper; all authors edited and approved the manuscript.

## Acknowledgements

We thank the Core Facility and the Cell Biology Facility of the Center of Biomedical Analysis, Tsinghua University for assistance with imaging. We thank Jiaying Yang for providing extracted total RNA from HeLa cells. This work was supported by funds from the Ministry of Science and Technology of China (grant number: 2016YFA0501100), the National Natural Science Foundation of China (grant numbers: 31925023, 31861143027, 21827810, and 31470721), the Junior Thousand Talents Program of China, Beijing Frontier Research Center for Biological Structure and Beijing Advanced Innovation Center for Structural Biology to Y.X. and the Key R & D Project of Shandong Province (grant number: 2019GSF108212) to D.Z.

